# A Web-resource for Nutrient Use Efficiency related Genes, QTLs, and microRNA in important cereals and model plants

**DOI:** 10.1101/222992

**Authors:** Anuj Kumar, Ajay Pandeya, Girik Malik, Kumari P Hima, Kumar S Anil, Mansi Sharma, Vijay Gahlaut, MNV Prasad Gajula, Prashanth Suravajhala, Krishna Pal Singh, Harindra Singh Balyan, Pushpendra K Gupta

## Abstract

Cereals are the key contributors to global food security. Genes involved in uptake (transport), assimilation and utilization of macro- and micro-nutrients are responsible for their content in grain and straw. Although many cereal genomic databases are available, currently there is no cohesive web-resource of manually curated nutrient use efficiency (NtUE) related genes and QTLs, etc. In this study, we present a web-resource containing information on NtUE related genes/QTLs and the corresponding available microRNAs for some of these genes in four major cereal crops [wheat (*Triticum aestivum*), rice (*Oryza sativa*), maize (*Zea mays*), barley (*Hordeum vulgare*)], two alien species (*Triticum urartu* and *Aegilops tauschii*) related to wheat, and two model species including *Brachypodium distachyon* and *Arabidopsis thaliana*. Gene annotations integrated in the current web-resource were collected from the existing databases and the available literature. The primary goal of developing this web-resource is to provide descriptions of the NtUE related genes and their functional annotation. MicroRNA targeting some of the NtUE related genes and the quantitative trait loci (QTLs) for NtUE related traits are also included. The available information in the web-resource should help the users to readily search the desired information.

**Web-resource URL:** http://bioclues.org/NtUE/

## INTRODUCTION

The world population is expected to be >9.6 billion by 2050 (http://www.fao.org/livestock-environment/en/). As a consequence, the food production must increase by 70% to meet the growing demand for food; this translates into 3 billion tons of additional cereal grain (http://www.fao.org/fileadmin/templates/wsfs/docs/Issues_papers/HLEF2050GlobalAgriculture.pdf;). In view of this, the food and nutritional security have been central to the worldwide discussions leading to development of strategies for enhanced and sustainable food production (http://www.fao.org/news/story/en/item/35571/icode/). These strategies include development of cultivars that would be (i) resilient to climate change, (ii) tolerant to biotic and abiotic stresses and (iii) less demanding for fertilizers and water. In connection with the last strategy, it is well known that the present day high yielding cultivars of a majority of crops have high demand for inputs like water and nutrients.

Out of the 17 elements needed for plant growth and development, three non-mineral elements (C, H, O) are derived from CO_2_ and H_2_O and the remaining 14 mineral elements are ordinarily derived from the soil as inorganic salts. These mineral elements include (i) six macronutrients including N (nitrogen), P (phosphorous), K (potassium), S (sulphur), Ca (calcium) and Mg (magnesium) that are required in large quantities (1000 to 15000 mg/kg of plant dry weight) and (ii) eight micronutrients including B (boron), Cl (chlorine), Cu (copper), Fe (iron), Mn (manganese), Mb (molybdenum) Zn (zinc) and Ni (nickel) that are needed in relatively small quantities (0.1 to 100 mg/kg of plant dry weight) [1].

Since most of the soils are deficient in N, P and K, these elements are applied to the crops in the form of chemical fertilizers to help their proper growth and development, eventually leading to improved grain yield [2, 3]. Among these three macronutrients, N and P are required in highest quantities by the plants 4. Further, with the increased future demand for cereal grains, especially within the developing countries, the demand for N and P is likely to grow 5.

It is also known that during the last four decades, the doubling of food production worldwide has been associated with 7-fold increase in consumption of N fertilizers and 3.5 fold increases in consumption of P fertilizers [6, 7]. It is estimated that further doubling of global food production during the next four decades would require a 3.15 fold increase in the total N application and 2.5 fold increase in total P application [6]. The enhanced use of chemical fertilizers would also add up to 50% (for N fertilizers) of the operational costs in agriculture [2].

It is estimated that 30 % to 50% of N applied in the fields is used by the crops [8], and the excess N and ammonia are converted by soil bacteria into nitrate and nitrite, which are lost *via* emission in the form of NH_3_, N_2_O, NO_2_ gases and/or leached out into the soil in the form of nitrate ions (NO^−^_3_) (2, 9), causing eutrophication of terrestrial and aquatic systems. In comparison to CO_2_, N_2_O is 300× more potent greenhouse gas (GHG) and 70% of this gas is contributed by natural sources causing a very high imbalance in GHGs, which play a significant role in climate change [5, 10].

Unlike N, a part of P derived from applied chemical fertilizers is held very tightly to the surface of the soil particles or is fixed as organic P compounds and therefore, remains unavailable to the plant for uptake; P is also lost through leaching into the ground and surface waters, damaging the surrounding environment [11]. The leached P also leads to eutrophication and associated algal blooms, decreased dissolved O2, foul odour, and in general, poor water quality [12]. To further exacerbate the problem, the P used in fertilizers is obtained from rock phosphate, which is a non-renewable source and could be depleted in 50-100 years [13]. Indirectly through the food chain, enhanced N and P fertilizer applications also leads to widespread multi-nutrient deficiencies in humans, thus becoming a health hazard.

In view of the importance of nutrient use efficiency (NtUE) as an agronomic practice in crop breeding, the European Commission recently concluded a five year research project ‘NUE-CROPS’ with the aim of improving NtUE in four major European food, feed and biofuel crops to reduce the negative environmental impact of crop production (EU-FP7 222-645; http://ec.europa.eu/research/fp7/pdf/19072010/nue_crops_-_kbbe.pdf). The findings of this project were discussed at a EUCARPIA section meeting on ‘Organic and Low-input Agriculture’ during 24-26, 2013 at the George August University of Gottingen and the proceedings were published in Euphytica (Volume 119, 2014).

It has been shown that traditional breeding practices involving higher application of chemical fertilizers would no longer provide yield improvements [14]. Both the FAO and UN have shown that ever since mid-1980s, the cereal crops including wheat, maize and also soybean have shown only 1% annual growth in their productivity, whereas in the developed countries the annual growth in crop yield has remained static with no improvement in productivity [15]. Thus the challenge before the geneticists and the plant breeders is to increase productivity of cereals in particular and that of the other food crops in general without compromising the quality of food and causing damage to the environment. Amongst the suggested strategies, a genetic intervention for the development of more NtUE crop plants is the most environment-friendly approach. This approach would require reduced fertilizer application associated with either an increased grain yield or with at least the current yield levels [5, 7, 11, 15]. Also termed as “An Evergreen Revolution” and ‘Second Green Revolution,’ this approach would allow increased productivity following sustainable agricultural practices [16, 17].

An increased understanding of the NtUE at the genetic and molecular levels would certainly help in developing NtUE cultivars giving enhanced crop productivity. The NtUE (ratio of grain yield and unit nutrient supplied or available) can be divided in two components: (i) nutrient uptake efficiency (NtptE = ratio of total plant nutrient and the available or supplied nutrient), which is the ability of the crop plant to extract nutrients from the soil, and (ii) nutrient utilization efficiency (NtutE = ratio of grain yield and total plant nutrients in grain and straw), which measures the capacity of the plant to convert the absorbed nutrient into grain yield [18], [19]. Another approach is to calculate the above ground biomass NtUE, which is measured by dividing the dry shoot weight by the nutrient supplied [20], but the plant breeders are most often interested in the first approach of measuring the NtUE in breeding of grain crops.

The NtUE is a complex trait involving signaling, acquisition, transport and utilization, the latter constituting assimilation and translocation/remobilization [21]. The detailed description of the mechanisms of signaling, uptake, transport and utilization of the nutrients is neither necessary, nor desirable in this article, but we know that they form an interwoven network of genetic control. Thus identification of genes/QTLs and the microRNA (miRNA) targeting the NtUE related genes involved in the uptake, transport and utilization of nutrients in different parts of the plant are important targets for breeding cultivars with improved NtUE. The genes that have been identified or are likely to be identified in future or the DNA markers associated with these genes/QTLs as well as the information related to miRNA can be exploited as follows: (i) marker-assisted selection (MAS) aimed at the selection of desirable genotypes exhibiting improved plant performance under reduced nutrient supply; (ii) a study of allelic variation for the genes/QTLs for their subsequent exploitation in plant breeding, and (iii) the development of transgenics with improved NtUE [21-23]. It may be noted that a number of genomic resources are freely available for cereals and include the following: Gramene [24], PIGD [25], PlantTFDB 3.0 [26], PSPDB [27], CerealsDB [28], MetaCrop 2.0 [29], Phytozome [30], PlantGDB [31], CSRDB [32], CR-EST [33], and GrainGenes [34]. However, similar web-resource for NtUE genes/QTLs, etc. is not available, although the USDA has developed a tool for calculating the approximate amounts of nutrients (N, P and K) required for an optimum harvest of individual agricultural crops (http://plants.usda.gov/npk/main). Therefore, we have developed NtUE web-resource (http://bioclues.org/NtUE/), which provides comprehensive information on genes related to the components of NtUE that are involved in uptake, transport, nitrate assimilation, reassimilation, amino acid biosynthesis, C/N storage and metabolism, signaling and regulation, translocation, remobilization, senescence, regulation, DNA-binding, ion-binding, copper homeostasis, etc. The web-resource also contains information on QTLs for different NtUE related traits like shoot growth, total N, nitrate, free-amino acid contents, relative grain yield, relative biomass yield, relative grain N, relative biomass N, N response, leaf length, P starvation, Fe and Zn homeostasis, etc. The miRNA targeting some of the NtUE related genes are also included in the web-resource. The present web-resource is first of the kind to the best of author’s knowledge. The authors believe that the web-resource would be useful for plant community, especially the plant breeders, who may like to plan experiments related to improvement of NtUE of cereal crop plants.

## METHODS AND WEB-RESOURCE CONTENTS

### Collection of NtUE responsive genes

We populated the entries in the web-resource by retrieving NtUE related genes using gene ontology (GO) keyword search, *viz*. nutrients, plants, crops and their NtUE related role from GenBank (http://www.ncbi.nlm.nih.gov/). EnsemblPlants (http://plants.ensembl.org/index.html), Gramene (http://www.gramene.org) and UniProt (http://www.uniprot.org/) [35]. Further, we also extensively searched PubMed with different keywords or their combination to find the relevant articles.

### Annotation of NtUE genes

We performed functional annotation studies at gene and protein level to develop comprehensive information for the collected NtUE responsive genes. Complementary and target-site accessibility both are important factors of plant regulatory small RNA target recognition mechanism [36]. To predict potential miRNAs targeting the NtUE responsive genes in plant species, the web-based psRNATarget server [37] was used with the default parameters. Further the QTLs for NtUE related traits were manually curated from the available scientific literatures, GrainGenes database (http://wheat.pw.usda.gov/GG3/) and MaizeGDB (http://www.maizegdb.org/).

### Database construction and implementation

The NtUE web-resource was setup based on three-tier architecture concept using Apache/PHP/MySQL on a window platform. An integrated system driven through MySQL (5.6.21), and PHP (5.6.24) was developed to handle the storage of annotated 688 genes in the web-resource. A flowchart depicting the steps involved in preparation of web-resource and acquisition of data by the user is presented in Figure 1.

**Figure 1.**
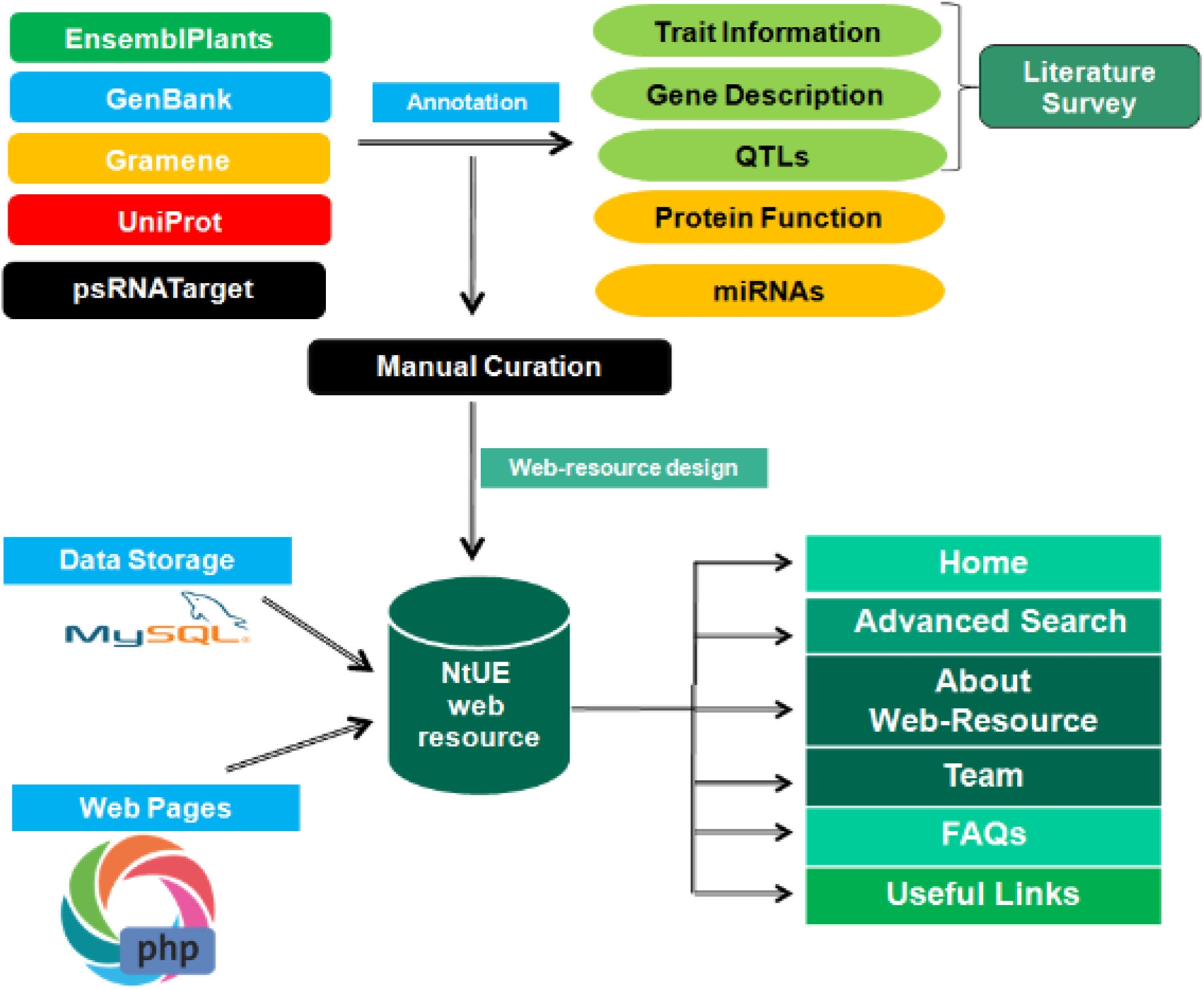
Flow chart depicting steps involved in preparation of NtUE web-resource and acquisition of the stored data.

## RESULTS AND DISCUSSION

NtUE is a comprehensive web-resource containing information on NtUE related genes/QTLs and the targeting microRNAs for some of these genes in three major cereal crops [wheat (*T. aestivum*), rice (O. *sativa*), maize (*Z. mays*)], two alien species (*T. urartu* and *A. tauschii*) related to wheat, and two model species including *B. distachyon* and *A. thaliana*.

In the current release, we have compiled 688 NtUE responsive gene records that were manually curated and annotated using different bioinformatics based methods. Each entry embedded in this web-resource has been arranged in gene centric manner for easy access and retrieval.

### Gene-Centric Portal

Each entry embedded in web-resource provides the following comprehensive information on NtUE responsive genes: (i) organism name; (ii) type of nutrients (macronutrients and micronutrients); (iii) genes symbol; (iv) gene ID (entry accession on UniProt, GenBank, Gramene and EnsemblPlants) with an active hyperlink which provides genomic information and sequences of entry on portal of parental database; (v) gene description; (vi) function; (vii) linked miRNAs with a active hyperlink to miRBase database; (viii) putative QTLs; (ix) chromosomal location marker; (x) experimental condition trait information; and (xi) references for QTLs as shown in (Figure 2).

**Figure 2.**
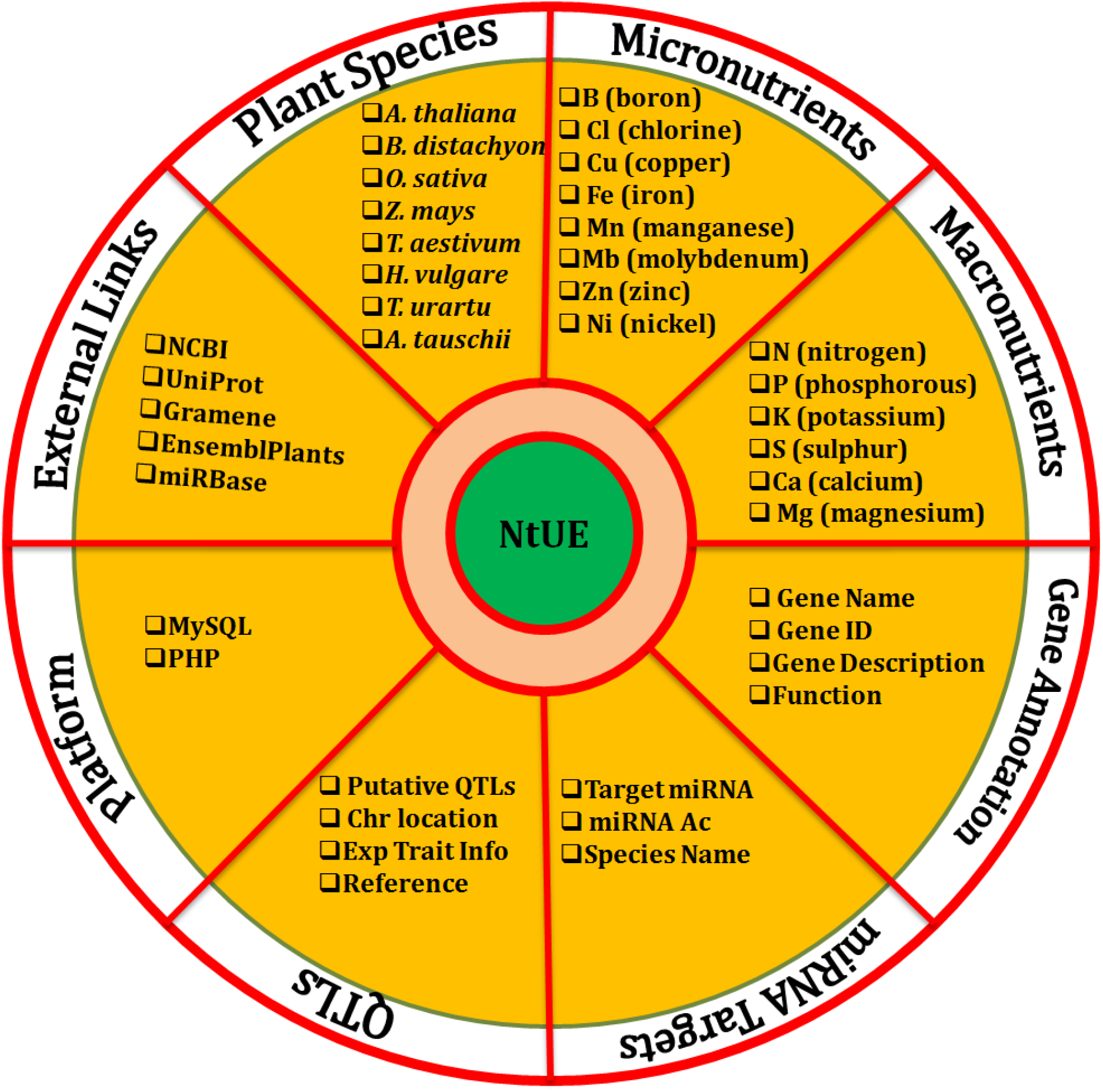
Schematic showing information available in NtUE web-resource.

### Data access

The information available in the web-resource could be accessed through keyword search tab by assigning the nutrients name (*i.e*. Nitrogen, Phosphorus, Potassium, Zinc, Iron etc.), organism name (*i.e. Arabidopsis thaliana, Brachypodium distachyon, Oryza sativa, Triticum aestivum, Hordeum vulgare*) etc.), gene name (*i.e*. NRT1.1, PHT1.1 etc.), accession number and clicking on the “search” button. After submitting the keyword, a page will open presenting a list of NtUE responsive genes details which contain the specific function. Further clicking on a ‘MORE INFORMATION’, a web page will direct the user to complete data details to particular gene. This scroll down based page provides comprehensive information that includes a summary of gene including: organism name, type of nutrients (macronutrients and micronutrients), genes information (gene symbol, gene ID, gene description, function) target miRNAs, and QTLs (chromosomal location marker, experimental condition trait information and references for QTLs). Alternatively, the user can search NtUE web-resource using advanced search option which will allow them to conduct organism specific search using gene name (*i.e* NRT1.1, PHT1.1 etc.) or accession number. Using a specific example of search option, the steps involved in searching keyword based search and advanced search options are shown in Figure 3. NtUE web-resource has a user-friendly entry point for each gene/miRNA/QTL. Each gene with corresponding miRNA and the record of the corresponding QTL reference article in web-resource is linkout to external resources/databases/journals like NCBI, UniProt, Gramene, EnsemblPlants, miRBase etc. Using external links, users can get detailed genomic and proteomic information of NtUE associated genes, miRNA target genes and the trait for each QTL. Additionally, it was ensured that standardized terminologies are used throughout, validated and processed making a good resource for interoperability of NtUE data. QTLs embedded in NtUE web-resource contain the chromosomal location, marker interval, trait information, and reference article with the external link to journal in which article published. Figure 3 shows an example of how more information on genes and targeted miRNAs can be retrieved.

**Figure 3.**
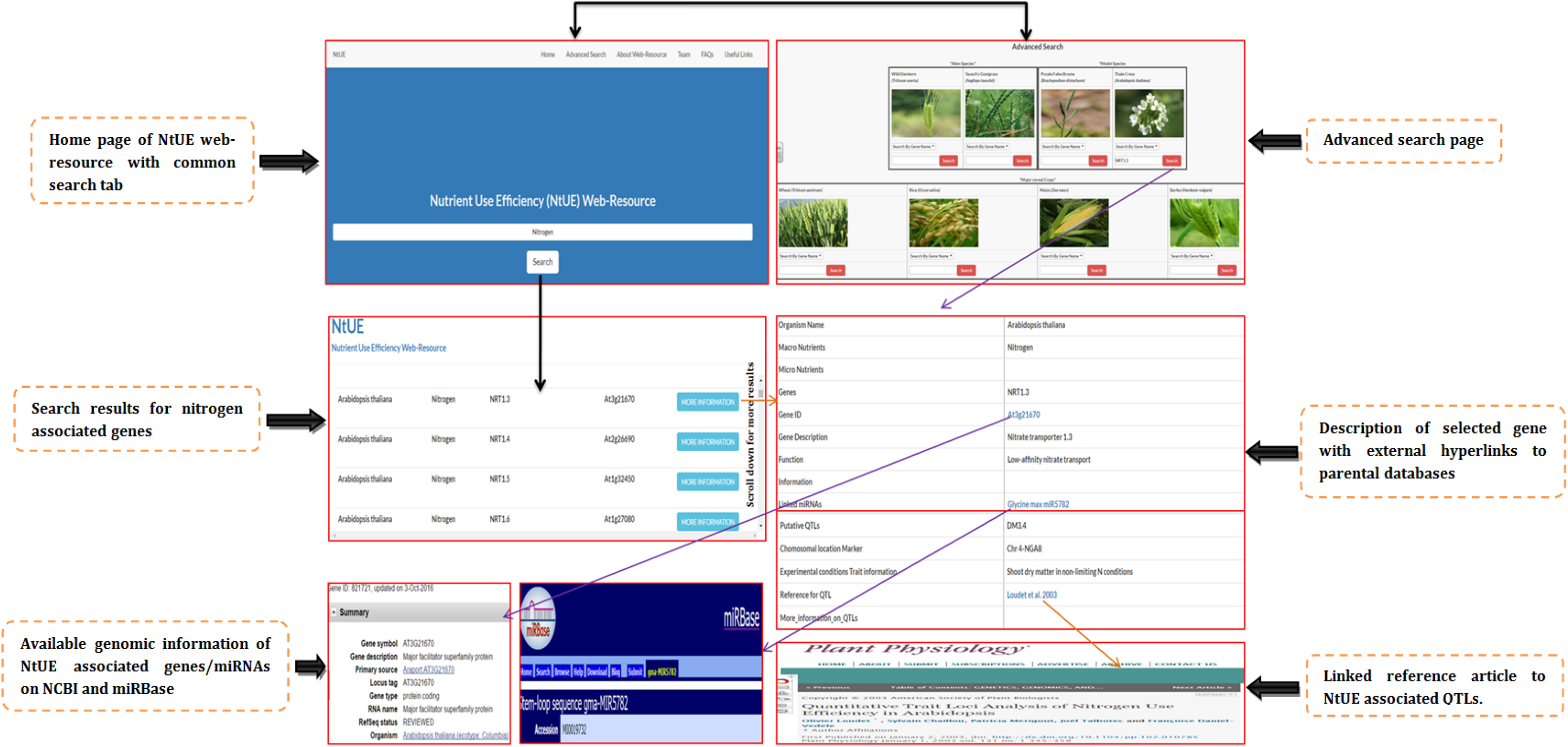
An illustration of the home page, search options, search summary and search results along with details of genomic information of NtUE associated gene on NCBI, information on miRBase database regarding targeted miRNA and linked reference article to QTLs on journal homepage.

## STATISTICS AND DATA ANALYSIS

### Type of nutrients

At present, NtUE web-resource covers 668 NtUE responsive genes corresponding to 12 nutrients types and the nutrient-wise distribution of NtUE genes in NtUE web-resource shown in the Figure 4. It is clear from the statistical graph that majority of NtUE genes are belongs to N (nitrogen). The other major nutrients embedded in web-resource are K (potassium), (P) phosphorus, Cu (copper), and Zn (zinc) in which NtUE genes have been detected.

**Figure 4.**
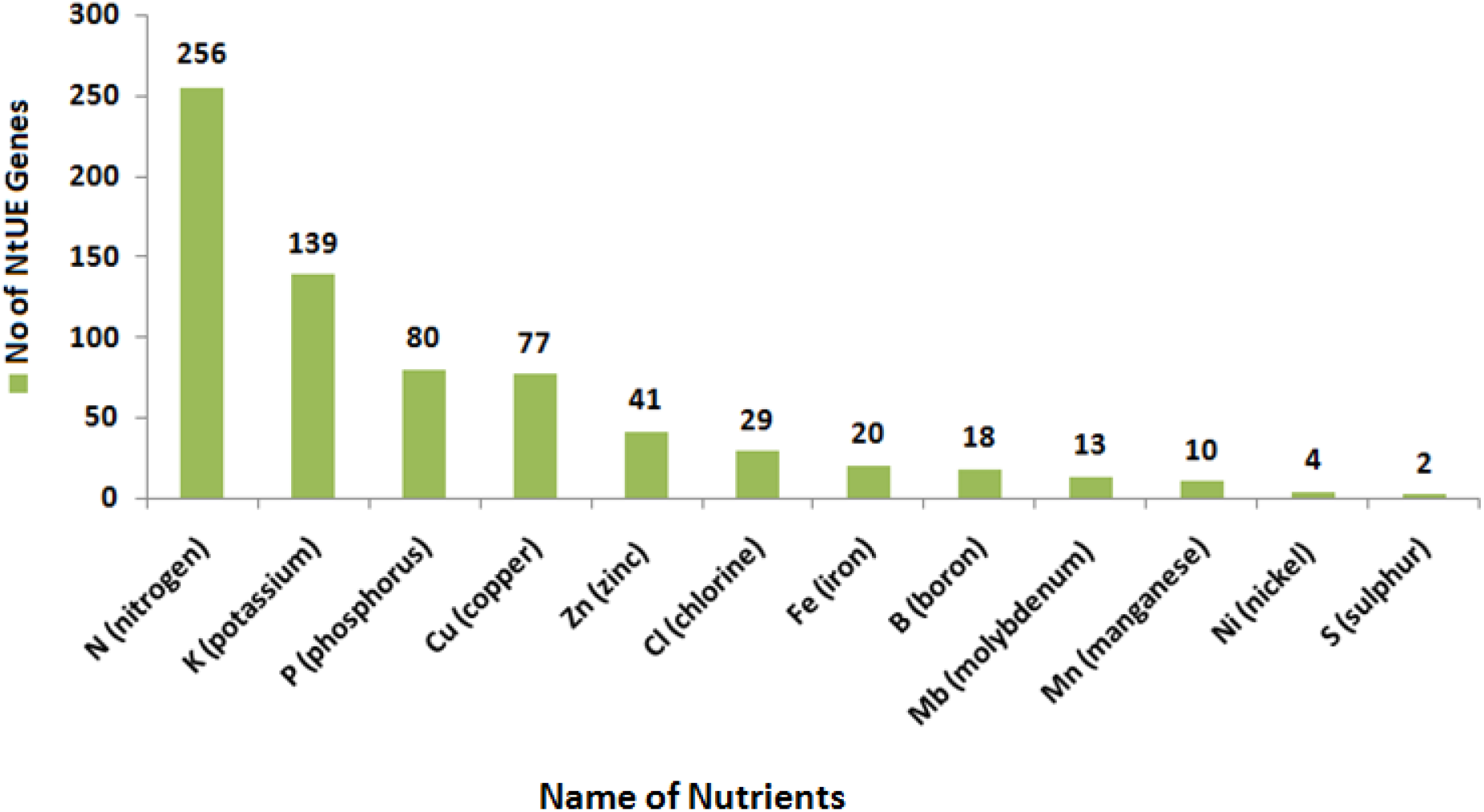
Distribution of macro and micronutrients responsive genes corresponding to their nutrient type in NtUE web-resource.

### NtUE responsive genes in cereal and model plant species

NtUE responsive genes have been reported in literature published in peer-reviewed journals to be detected from genomes of 8 model and cereal plant species (emebedded in web-resource) including Arabidopsis, rice, wheat, maize etc. (Figure 5). Among, all the embedded plant genomes Arabidopsis are the most explored source of annotated NtUE responsive genes in NtUE web-resource.

**Figure 5.**
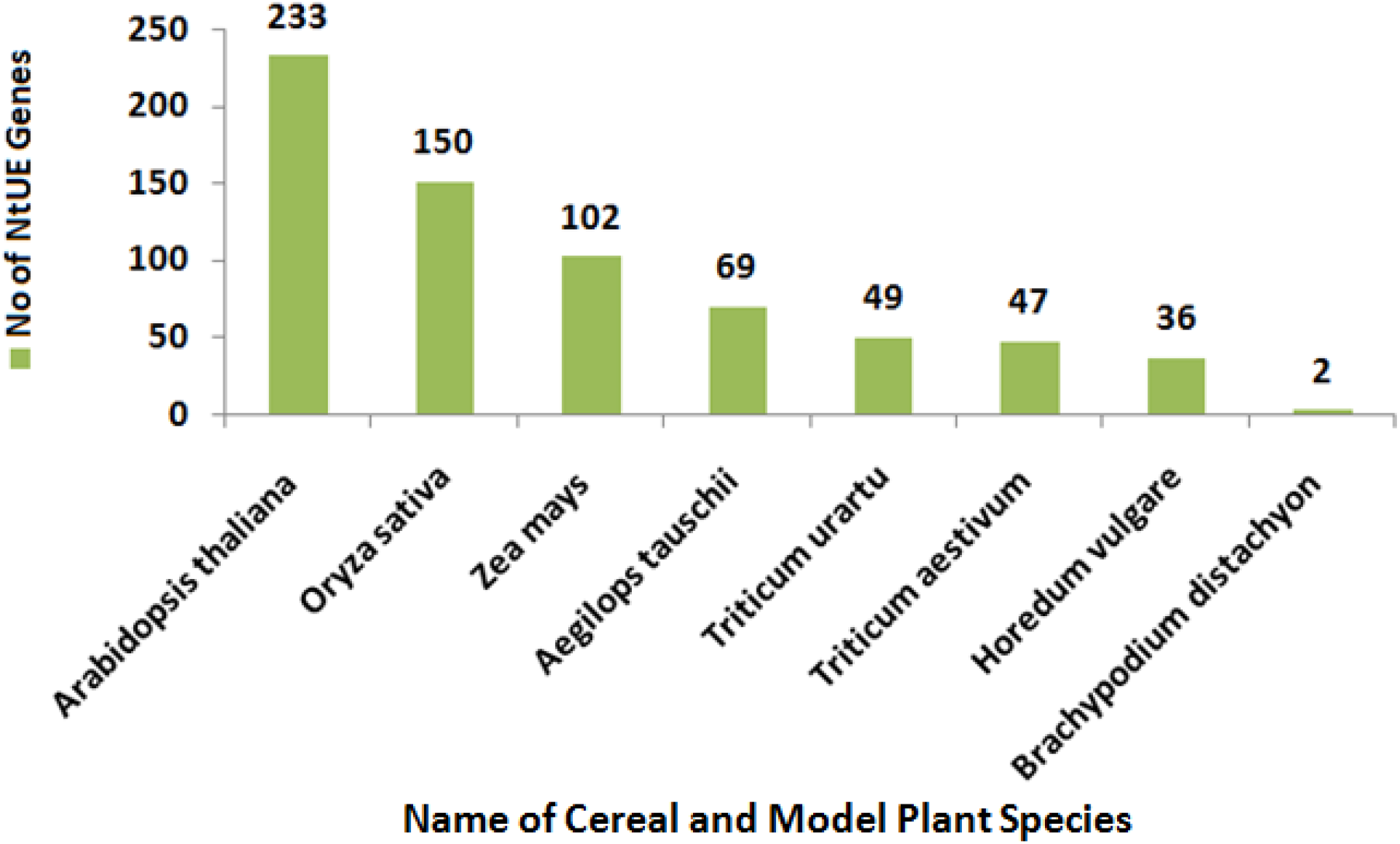
Distribution of macro and micronutrients responsive genes in cereal and model plant species embedded in NtUE web-resource.

### Comparison of NtUE web-resource with Gramene

Gramene is a biological repository which provides genomic information about members of poaceae family. It is a experimental based comprehensive database consist with genomic sequences, chromosomal location, associated QTLs and markers of genes from various popular genomes belongs to poaceae family. Whereas, NtUE web-resource is a literature curated resource for models and cereals NtUE responsive genes that exhibit variation in different nutrient and plant genomes. It is a unique resource that not only lists the NtUE genes according to their nutrient type but also provides data on various aspects like miRNA-targets, associated QTLs, marker information to detect QTLs mapping, trait information, experiment information, external link to QTL reference article etc. Further, on comparing NtUE web-resource with Gramene its uniqueness is established as we found that only NtUE genes form cereal are overlapping with Gramene while NtUE genes form model plant species are unique to NtUE web-resource.

## CONCLUSION AND FUTURE DIRECTION

NtUE web-resource is a first of its kind attempt to provide a comprehensive non-redundant catalogue of NtUE genes play a vital role in model and cereal plant species during plant growth and development, supporting evidence from published literature. We hope that this web-resource would be a very useful compendium to the geneticists, plant breeders and computational biologists involved in crop improvement. It contains manually curated entries with an adequate level of stringency to select only those genes/QTLs/miRNAs, which are validated and linked to NtUE. This resource will help in providing new solutions to plant geneticists and breeders, who are aiming at more sustainable use of limited soil nutrients for enhanced productivity without perhaps compromising the nutritional quality of the grains. Information on more than half of these genes, QTLs and miRNAs is not available in the previously reported databases. With further progress in the identification of genes/QTLs/miRNAs for NtUE related traits, we anticipate that these numbers will eventually grow by at least a two-to three-folds.

## Author contribution statement

PKG and HSB conceived, outlined, edited and wrote a part of the manuscript. AK and PS prepared outline and wrote the first draft of manuscript and assisted PKG and HSB in finalizing the manuscript. AP, GM and MNVP developed all APIs and web-resource search interfaces. AK, AKS, VG and HKP manually curated and annotated the web-resource entries. AK, KPS and PS manually cross-checked all the annotations. MS contributed in statistics and data analysis. All authors agreed and have gone through the final version of the manuscript.

## ACKNOWLEDGMENTS

This study was undertaken utilizing the DBT-sponsored Bioinformatics Infrastructure Facility (BIF) of the Department of Genetics and Plant Breeding, Ch. Charan Singh University, Meerut, India. PKG and HSB held the position of INSA-Senior Scientist during the period of study. PKG earlier held the position of Senior Scientist at National Academy of Sciences India (NASI), Allahabad. AK was awarded a JRF/SRF under the scheme of DBT-BTISnet program.VG was awarded a JRF under the Senior Scientist Platinum Jubilee Fellowship program, and was later awarded the position of SRF/RA under a DBT project. We thank Dr. VS Sundararajan for his useful suggestions on the prototype of the database. We also thankful to Prof. PB Kavi Kishor to proof read the MS.

